# PolyQ expansion does not alter the Huntingtin-HAP40 complex

**DOI:** 10.1101/2021.02.02.429316

**Authors:** Bin Huang, Qiang Guo, Marie L. Niedermeier, Jingdong Cheng, Tatjana Engler, Melanie Maurer, Alexander Pautsch, Wolfgang Baumeister, Florian Stengel, Stefan Kochanek, Rubén Fernández-Busnadiego

**Affiliations:** Department of Gene Therapy, Ulm University, 89081 Ulm, Germany; Department of Molecular Structural Biology, Max Planck Institute of Biochemistry, 82152 Martinsried, Germany; State Key Laboratory of Protein and Plant Gene Research, School of Life Sciences and Peking-Tsinghua Center for Life Sciences, Peking University, 100871Beijing, China; Department of Biology, University of Konstanz, 78457 Konstanz, Germany; Konstanz Research School Chemical Biology, University of Konstanz; Gene Center, Department of Biochemistry and Center for integrated Protein Science Munich, Ludwig-Maximilians University, 81377 Munich, Germany; Department of Medicinal Chemistry, Boehringer Ingelheim Pharma GmbH & Co. KG, 88397 Biberach an der Riß, Germany; Institute of Neuropathology, University Medical Center Göttingen, 37099 Göttingen, Germany; Cluster of Excellence “Multiscale Bioimaging: from Molecular Machines to Networks of Excitable Cells” (MBExC), University of Göttingen, 37075 Göttingen, Germany

**Keywords:** Huntingtin, Huntington’s disease, polyglutamine expansion, cryo-electron microscopy, crosslinking mass spectrometry

## Abstract

The abnormal amplification of a CAG repeat in the gene coding for huntingtin (HTT) leads to Huntington disease (HD). At the protein level, this translates into the expansion of a poly-glutamine (polyQ) stretch located at the HTT N-terminus, which renders it aggregation-prone by unknown mechanisms. Here we investigated the effects of polyQ expansion on HTT in a complex with its stabilizing interaction partner huntingtin-associated protein 40 (HAP40). Surprisingly, our comprehensive biophysical, crosslinking mass spectrometry and cryo-EM experiments revealed no major differences in the conformation of HTT-HAP40 complexes of various polyQ length, including 17QHTT-HAP40 (wild type), 46QHTT-HAP40 (typical polyQ length in HD patients) and 128QHTT-HAP40 (extreme polyQ length). Thus, HTT polyQ expansion does not alter the global structure of HTT when associated with HAP40.

## Introduction

Huntingtin (HTT) is a large protein (348 kDa) involved in a wide variety of cellular processes, including transcriptional regulation, vesicular transport, endocytosis and autophagy (Saudou and Humbert, 2016). The expansion of a CAG repeat in the first exon of the *HTT* gene translates into an expanded poly-glutamine (polyQ) repeat at the N-terminus of HTT and causes Huntington’s disease (HD). HD is a devastating neurodegenerative syndrome characterized by uncontrolled movements (chorea), emotional instability and cognitive impairment (Bates et al., 2015; Finkbeiner, 2011). The age of HD onset correlates with the length of HTT polyQ repeat, suggesting an important pathological role of this amino acid stretch. Furthermore, polyQ expansion renders N-terminal fragments of HTT aggregation-prone *in vitro* and *in vivo* (Scherzinger et al., 1997), and similar fragments form inclusion bodies in the brain of HD patient (Difiglia et al., 1995). The effects of polyQ expansion on full length HTT are less well understood, but recent studies reported some conformational changes (Jung et al., 2020; Vijayvargia et al., 2016). However, only low-resolution structures of polyQ-expanded HTT were reported, limiting detailed understanding of the effects of polyQ expansion.

We have recently shown that the formation of a complex with huntingtin-associated protein 40 (HAP40), a very abundant cellular binding partner of HTT, dramatically reduces the structural flexibility of HTT, allowing structure determination by cryo-electron microscopy (cryo-EM) (Guo et al., 2018). The formation of this complex may be implicated in the recruitment of HTT to Rab5-positive endosomes (Pal et al., 2006). The HTT-HAP40 complex structure revealed that HTT is a largely alpha-helical protein, with most helices arranged in tandem repeats (Guo et al., 2018). HTT is divided in three major domains. The N- and C-terminal domains (“N-HEAT” and “C-HEAT”) are rich in HEAT (huntingtin, elongation factor 3, protein phosphatase 2A and lipid kinase TOR) repeats arranged in a solenoid fashion. These domains are connected by a smaller bridge domain. HAP40 is also largely α-helical and has mainly a tetratricopeptide repeat (TPR)-like organization. The cryo-EM structure revealed only weak connections between the domains, explaining HTT’s conformational flexibility. In the complex with HAP40 this flexibility is counteracted by HAP40 binding within a cleft, contacting the N-HEAT and C-HEAT domains mainly by hydrophobic interactions and the bridge domain by electrostatic interactions. Besides these domains, several presumably unfolded regions, including the polyQ stretch, were not resolved in the structure.

Here we used the HTT-HAP40 complex as a tool to examine the effects of pathological polyQ expansion on HTT structure. We cloned and expressed three complexes: 17QHTT-HAP40 (WT), 46QHTT-HAP40 (typical polyQ length in HD patients) and 128QHTT-HAP40 (extreme polyQ length). Combining biophysical techniques, cross-linking mass spectrometry and high-resolution structural determination by cryo-EM, we conclude that the polyQ length does not substantially influence the global architecture of the HTT-HAP40 complex.

## Results

### Purification, biochemical and biophysical characterization of HTT-HAP40

Previously, we described the production of human full length 17QHTT as a complex with HAP40 in a human cell line (Guo et al., 2018). To also enable the production and analysis of polyQ-expanded HTT, we generated additional human cell lines expressing full length HTT containing either 46 (46QHTT) or 128 glutamines (128QHTT). In these cells, HTT was expressed after induction with doxycycline, while HAP40 was under constitutive promoter control. We used a C-terminal Strep-tag on HAP40 for affinity purification of the HTT-HAP40 complexes, followed by size-exclusion chromatography (SEC).

Independent of their polyQ length, all HTT-HAP40 complexes eluted in a practically identical manner as a symmetric narrow peak and in similar yields (Figure 1A). Coomassie staining and Western blot analyses confirmed the purity of the materials, the presence of HTT and HAP40 in the complexes and the differences in polyQ length (Figure 1B, C). Both differential scanning fluorimetry (Figure 1D) and differential intrinsic tryptophan scanning fluorimetry (Figure 1E) showed very similar and sharp unfolding profiles and nearly identical melting points, indicating that the stabilities of the three complexes were identical. Altogether, these data show that HTT polyQ expansion does not induce major changes in the shape, stability and aggregation propensity of HTT-HAP40 complexes.

**Figure 1:**
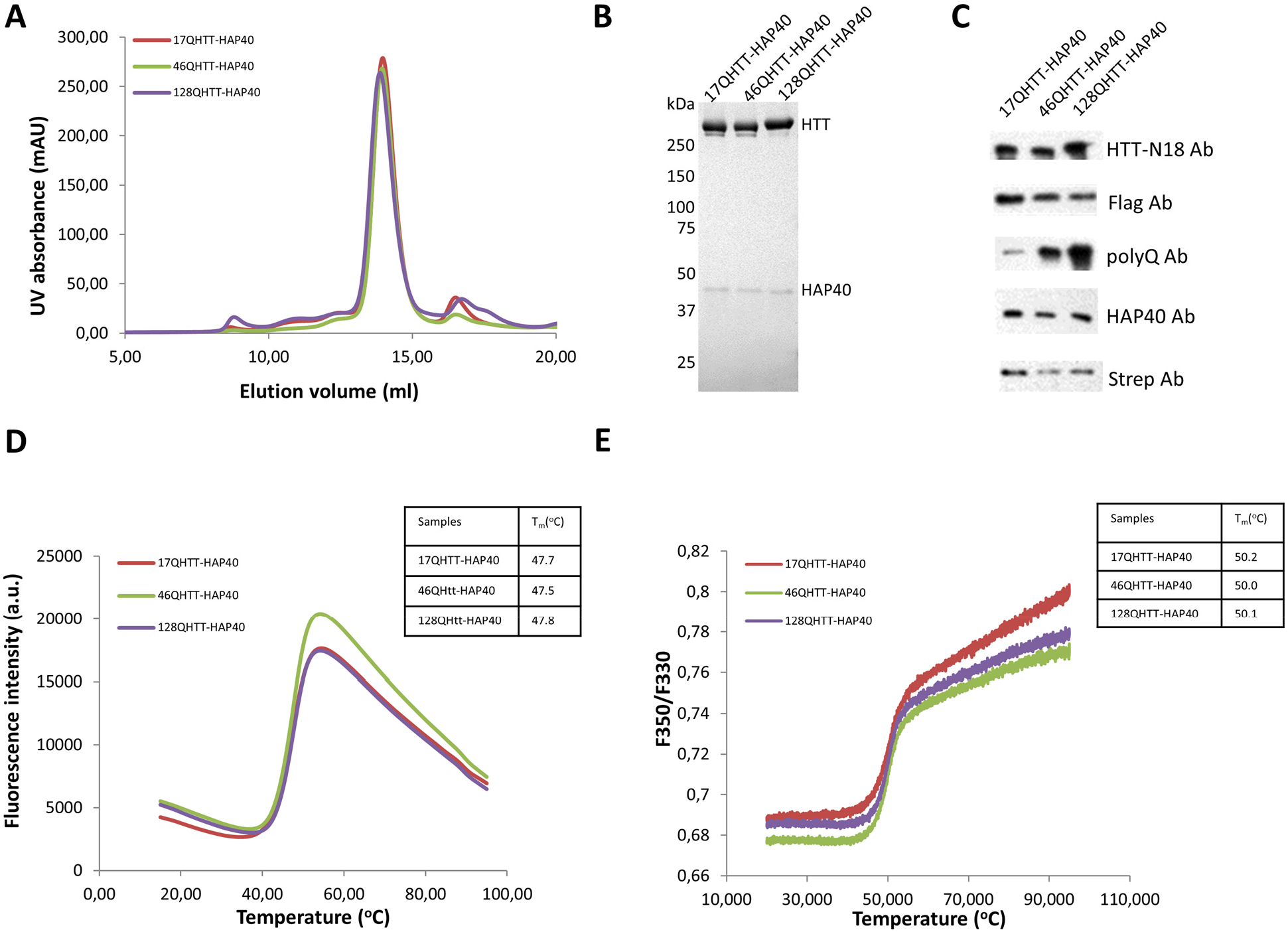
Purification, biochemical and biophysical characterization of HTT-HAP40 complexes. **(A)** SEC elution profiles of HTT-HAP40 complexes differing in HTT polyQ length. The HTT-HAP40 complexes were purified from HEK293-based human cells co-expressing FLAG-tagged HTT with 17Q (17QHTT), 46Q (46QHTT) or 128Q (128QHTT), respectively, together with Strep-tagged HAP40. Cleared cell lysates were incubated with Strep-Tactin beads, washed and bound proteins were eluted with desthiobiotin prior to loading on a Superose 6 10/300 increase column. **(B)** Coomassie staining of the main peak of the different HTT-HAP40 eluates shown in (A). **(C)** Western Blot analyses of HTT-HAP40 eluates after affinity purification and SEC, using antibodies detecting HTT (upper two rows), polyQ (middle) and HAP40 (lower two rows). **(D, E)** Thermal unfolding of the different HTT-HAP40 complexes. Melting curves and melting points were obtained by differential scanning fluorimetry (DSF) (D) and nano differential scanning fluorimetry (nanoDSF) (E). Melting temperatures are indicated. One of three measurements is shown.

### Cryo-EM analysis of polyQ-expanded HTT-HAP40 complexes

To further investigate possible structural rearrangements, we analyzed polyQ-expanded HTT-HAP40 complexes by cryo-EM (Figure 2). The structures of 46QHTT-HAP40 (Figure 2A) and 128QHTT-HAP40 (Figure 2C) were determined with global resolutions of 3.6 Å and 4.1 Å respectively (Figure S 1, Table S 1). No density was observed for the disordered insertions that were absent from the 17QHTT-HAP40 structure, including the polyQ stretch. Based on these maps, atomic models were built using energy minimization. Consistent with our biophysical data, the overall architectures of the 17QHTT-HAP40 (Guo et al., 2018) (PDB-6EZ8), 46QHTT-HAP40 and 128QHTT-HAP40 complexes were identical: two HEAT repeats domains linked by a bridge domain, with HAP40 binding in the cleft between the HTT domains. For a more detailed comparison, we calculated the root-mean-square deviation (RMSD) of Cα between the WT and polyQ-expanded atomic models (Figure 2B, D). The average RMSD were 0.89 Å (17Q vs 46Q) and 1.29 Å (17Q vs 128Q) respectively. The maximum RMSD was smaller than 3 Å in both cases, indicating that the structures are identical at the current resolution. Therefore, our cryo-EM data conclusively show that polyQ expansion has no noticeable influence on the architecture of the HTT-HAP40 complex.

**Figure 2:**
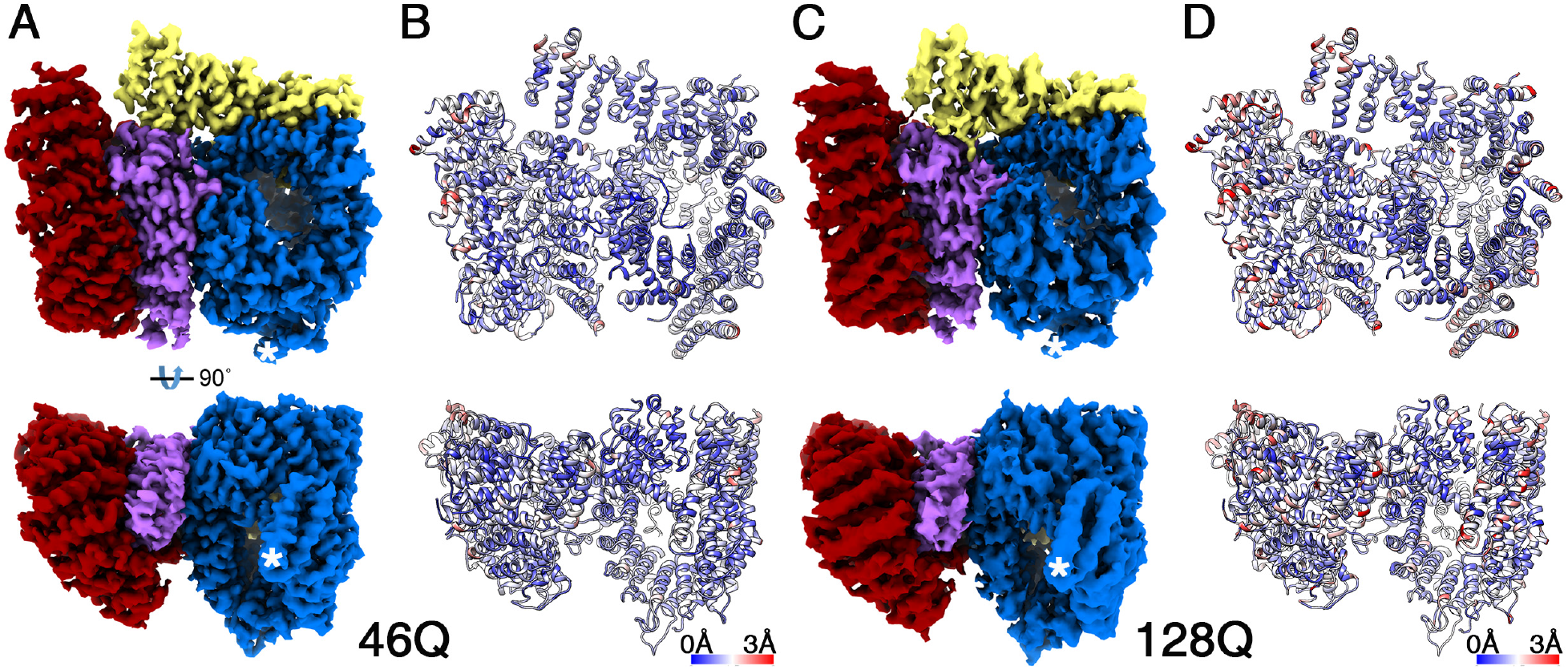
Architecture of polyQ expanded HTT-HAP40 complexes by cryo-EM. **(A, C)** Reconstructed density maps of 46QHTT-HAP40 (A) and 128QHTT-HAP40 (C) filtered according to local resolution and shown as surface representation. The HTT N-HEAT domain, bridge domain and C-HEAT domains are respectively colored in blue, yellow and maroon. HAP40 is colored in purple. Stars indicate the most N-terminal region of HTT resolved. **(B, D)** Atomic models of 46QHTT-HAP40 (B) and 128QHTT-HAP40 (D) are shown in ribbon representation, colored according to their RMSD with 17QHTT-HAP40 (Guo et al., 2018) (PDB-6ZE8). See also Figure S 1 and Table S 1.

### Crosslinking mass spectrometry

We reasoned that WT and polyQ-expanded HTT-HAP40 complexes could also differ in unstructured regions not resolved in the cryo-EM structures. We addressed this possibility using chemical crosslinking coupled to mass spectrometry (XL-MS). Isotopically labelled disuccinimidyl suberate (DSS) was used to crosslink lysine residues in 17QHTT-HAP40, 46QHTT-HAP40 and 128QHTT-HAP40 complexes. XL-MS uses covalent bonds formed by crosslinking reagents in order to identify crosslinking sites by MS that reflect the spatial proximity of regions within a given protein (intralink) or between different proteins or subunits in a protein complex (interlink). Additional information on the accessibility of a specific lysine residue can be obtained from crosslinks that react on one side with the protein and hydrolyse on the other side (monolink). Approximately 65 % of the 123 lysines present in HTT were accessible to DSS (77 lysines in 17QHTT, 75 in 46QHTT and 88 in 128QHTT; Table S 2). In all complexes, 52 lysines encompassing the whole length of HTT were modified by monolinks, while five monolinks were identified in at least two of the complexes and 14 were unique to a single complex (Figure S3 and Table S 2). Unique monolinks did not show any specific localization, suggesting that they were due to experimental variabilities.

We further identified 78 intra-HTT crosslinks, of which 49% were present in all three complexes (Figure 3, Figure S3 and Table S 2). Additionally, 14 crosslinks were common to two complexes and 26 were found only in one (Figure 3, Figure S3 and Table S 2). Many of the crosslinks involved unstructured regions not resolved in the cryo-EM structures, but no crosslinks were detected within HTT exon 1 (Figure S 2). In general, the crosslinks built up four main interaction clusters within HTT (Figure 3, Figure S3): i) N-HEAT domain, ii) N-HEAT to bridge domain, iii) N-HEAT to C-HEAT domain and iv) C-HEAT domain. The intra-domain (N-HEAT and C-HEAT) crosslinks were extremely similar for all complexes, while the inter-domain connectivity (N-HEAT to bridge and C-HEAT domains) was overall highly similar as well, but exhibited some small differences. However, as slight variations in the detected crosslinks are common even between replicate measurements (Table S2), our data strongly indicates no significant differences between the overall HTT crosslinking pattern (intra- and monolinks) of the different HTT-HAP40 complexes (Figure S3).

**Figure 3:**
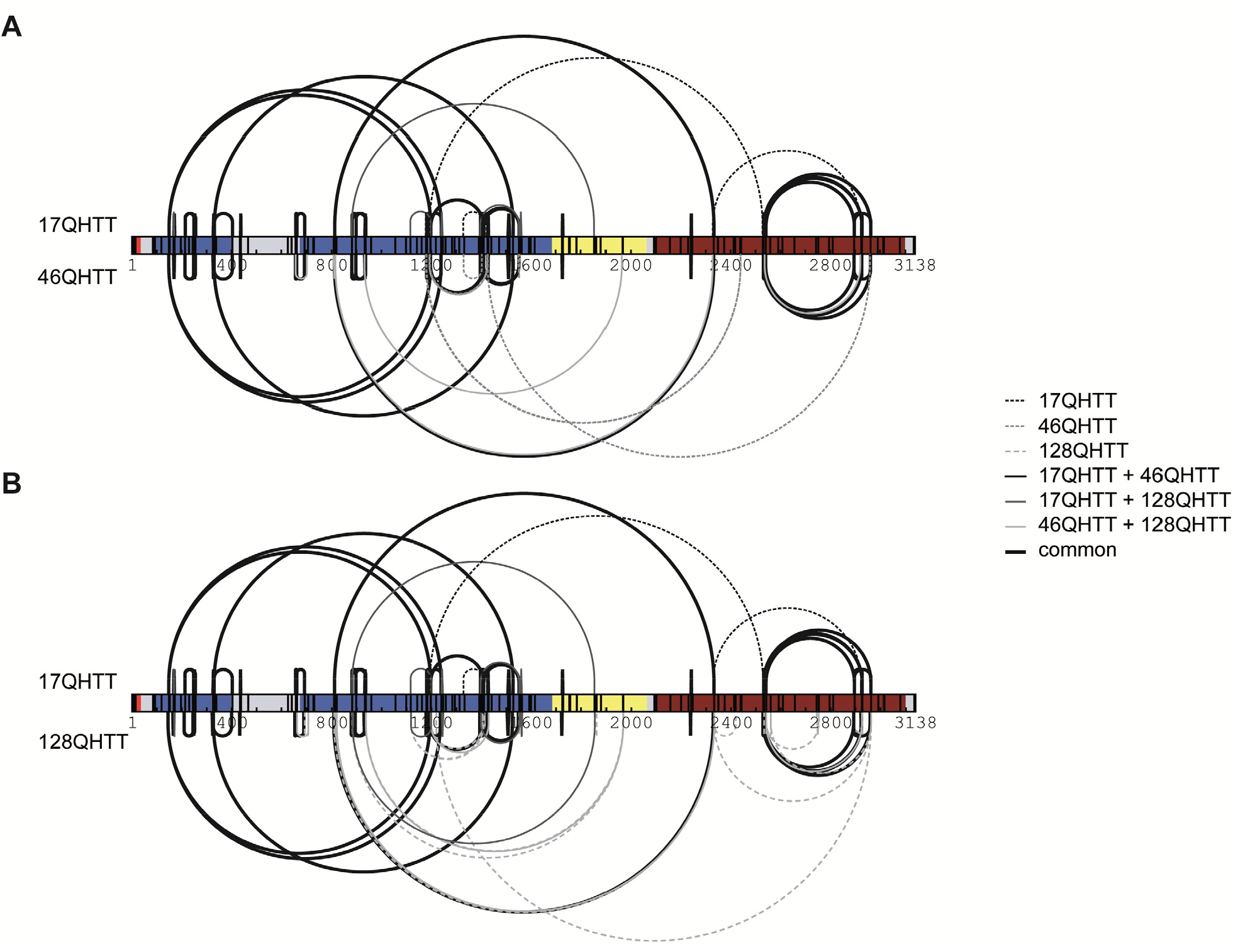
Effects of polyQ expansion on intra-HTT crosslink patterns of HTT-HAP40 complexes. **(A, B)** Comparative representation of all intra-HTT crosslinks identified within three different HTT-HAP40 complexes, namely 17QHTT-HAP40 vs 46QHTT-HAP40 (A) and 17QHTT-HAP40 vs 128QHTT-HAP40 (B). A uniform numbering scheme based on the 17QHTT sequence was used. Crosslinks common to all three complexes are represented by thick, black solid lines. Crosslinks identified in only two of the complexes are represented by thin solid lines (17QHTT and 46QHTT, black; 17QHTT and 128QHTT dark grey; 46QHTT and 128QHTT, light grey). Crosslinks identified in only one of the complexes are depicted by dotted lines (17QHTT, black; 46QHTT, dark grey; 128QHTT light grey). Only high-confidence crosslinks that were reliably identified at least in two out of three biological replicates are shown (see methods for details). The HTT N-HEAT domain, bridge domain and C-HEAT domains are colored in blue, yellow and maroon, respectively. HAP40 is colored in purple. See also Figure S 2, Figure S 3 and Table S 2.

In terms of inter-protein interactions (i.e. crosslinks between lysines within HTT and HAP40), 13 unique crosslinks were identified between HTT and HAP40, of which 9 were common for all three complexes, three were identified in two different complexes and one was uniquely identified in 46QHTT-HAP40 (Figure S 3 and Table S 2). Again, such a high overlap between the detected HTT-HAP40 crosslinks strongly indicates that the overall interaction pattern of the two proteins is not affected by HTT polyQ length (Figure S3). As residues crosslinked by DSS can be up to ∼30 Å apart (Iacobucci et al., 2019), all detected intra-HTT and inter-HTT-HAP40 crosslinks were fully compatible with our cryo-EM structures, reinforcing the consistency of our results.

Taken together, our biophysical, cryo-EM and crosslinking mass spectrometry data converge to rule out a major influence of HTT polyQ length on the architecture of the HTT-HAP40 complex.

## Discussion

The biophysical mechanisms by which polyQ expansion leads to HTT aggregation in HD remain poorly understood. Previous studies suggested that HTT could adopt different 3D conformations, some of which would be influenced by polyQ expansion (Pardo et al., 2010; Saudou and Humbert, 2016; Vijayvargia et al., 2016). At the same time, polyQ expansion does not result in an abrupt conformational change of the HTT N-terminus (Bravo-Arredondo et al., 2018; Newcombe et al., 2018; Warner et al., 2017). A recent study showed cryo-EM analyses at ∼10 Å resolution of recombinant 23QHTT and 78QHTT upon GraFix stabilization, in combination with XL-MS, small-angle X-ray scattering and hydrogen-deuterium exchange (Jung et al., 2020). This study indicated that apo HTT (i.e. HTT not bound to HAP40), has a flexible and modular character. While no significant differences within the major HTT domains were detected in 23QHTT and 78QHTT compared to the 17QHTT-HAP40 structure (Guo et al., 2018), changes in the relative positions between HTT domains resulted in a more extended conformation of 23QHTT and in particular 78QHTT.

In contrast, our high-resolution cryo-EM structure analyses show no differences in the overall architecture of the HTT-HAP40 complex irrespective of polyQ length. Although the HTT N-terminal exon 1 fragment may be more compacted with increasing polyQ length (Bravo-Arredondo et al., 2018; Newcombe et al., 2018; Warner et al., 2017), it was not visible in any of the structures, possibly due to the flexibility of the proline-rich region. These data also imply that the exon 1 fragment has no scaffolding function within the complex core, rationalizing why polyQ length does not affect the overall HTT-HAP40 structure.

Our structural view that HAP40 shields hydrophobic patches on opposite faces of the N-HEAT and C-HEAT domains independently of the polyQ length is consistent with our DSF results, thermal unfolding behavior and crosslink mass spectrometry. Furthermore, a large proportion of the intra-HTT crosslinks detected in our samples were also found in apo HTT with wild type and expanded polyQ lengths (Vijayvargia et al., 2016). Therefore, while polyQ expansion may influence the conformation of apo HTT, the stabilization of HTT conformation by HAP40 binding (Guo et al., 2018) is independent of polyQ length.

The increased flexibility of apo HTT may also explain its elution characteristics upon SEC and its sedimentation behavior in analytical ultracentrifugation (Huang et al., 2015). Those experiments revealed that a significant proportion of wild type HTT with normal polyQ length formed dimers, higher-order oligomers and aggregates. Under these *in vitro* conditions, oligomer and aggregate formation was further enhanced by increasing polyQ length (Huang et al., 2015). However, very little oligomer and aggregate formation was observed for 46QHTT and 128QHTT when purified as complex with HAP40. Thus, HTT-HAP40 complexes appear highly stable independent of HTT polyQ length.

Although HAP40 is an abundant interactor of HTT (Guo et al., 2018; Peters and Ross, 2001), there is little experimental information on HAP40 function(s). The HTT-HAP40 complex may be involved in endosomal function (Pal et al., 2006). We recently reported that HAP40 and HTT likely co-evolved in unikonts and that the HTT-HAP40 interaction is evolutionarily conserved, suggesting its functional importance (Seefelder et al., 2020). We also observed that HAP40 steady-state protein levels are directly dependent on HTT (both wild type and mutant), and that the otherwise short half-life of HAP40 is strongly increased by the interaction with HTT (Huang et al., submitted). Since it was not possible to reconstitute the HTT-HAP40 complex from purified monomers *in vitro* (Guo et al., 2018), we hypothesize that the complex is established in cells early after translation or even co-translationally, resulting in a rather rigid structure (Guo et al., 2018 and this report).

Altogether, we show that HTT polyQ expansion does not alter HTT structure within the HTT-HAP40 complex. Apo HTT, i.e. not bound to HAP40, may be more flexible and undergo subtle conformational changes upon polyQ expansion (Jung et al., 2020). Whether such rearrangements are related to the described loss-or gain-of-function phenotypes of mutant full length HTT in HD (Cisbani and Cicchetti, 2012; Zuccato et al., 2010) requires further investigation.

## Acknowledgments

We thank Günter Pfeifer and Jürgen Plitzko for electron microscopy support, Florian Beck for help with image processing. This work has been funded by the CHDI Foundation, the Huntington-Stiftung, the Max Planck Society, the European Commission (grant FP7 GA ERC-2012-SyG_318987–ToPAG to W.B.), the Konstanz Research School Chemical Biology and the Deutsche Forschungsgemeinschaft (DFG, German Research Foundation – project number 412854449 to S.K., Emmy Noether grant STE 2517/1-1 to F.S. and Germany’s Excellence Strategy grant EXC 2067/1-390729940 to R.F.-B.). M.N. is grateful for funding from the Zukunftskolleg.

## Author Contributions

B.H., S.K., M.N., F.S., A.P., W.B., Q.G., R.F.-B designed experiments. M.M., A.P., H.B. performed DSF and nanoDSF studies. M.N. performed crosslinking studies. H.B. prepared HTT-HAP40 samples for cryo-EM and performed biochemical analyses together with T.E. Q.G. performed cryo-EM work. J.C. built the atomic model. Q.G., J.C. and R.F.-B. analyzed the structure. B.H., S.K., M.N., F.S., Q.G., J.C., A.P., R.F.-B. analyzed data. B.H., Q.G., M.N. prepared figures. B.H., S.K., M.N., F.S., A.P., W.B., Q.G., R.F.-B wrote the manuscript. All authors commented on the manuscript.

## STAR Methods

### RESOURCE AVAILABILITY

#### Lead contact

Requests for resources and reagents should be addressed to the lead contact Prof. Rubén Fernández-Busnadiego

#### Materials Availability

Plasmids and cell lines generated in this study will be provided upon request.

#### Data and Code Availability

The cryo-EM maps of 46QHTT-HAP40 and 128QHTT-HAP40 have been deposited at the Electron Microscopy Data Bank under accession codes EMD-30911 and EMD-30912, respectively. The modeled structures of the 46QHTT-HAP40 complex and 46QHTT-HAP40 have been deposited at the Protein Data Bank under accession codes 7DXJ and 7DXK. The cryo-EM maps of 17QHTT-HAP40 have previously been deposited under accession codes EMD-3984 and PDB-6EZ8, respectively.

The MS raw files, databases containing protein fasta sequences for analysis with xQuest as well as xQuest result files have been deposited to the ProteomeXchange Consortium via the PRIDE (Vizcaino et al., 2016) partner repository with the dataset accession number PXD018451 (username; password: fnbVMbiT).

## EXPERIMENTAL MODEL AND SUBJECT DETAILS

Human HEK293-based cell lines for production of HTT-HAP40 complexes were propagated in MEM Alpha medium (GIBCO, #22561-021), supplemented with 10 % fetal bovine serum (GIBCO, #10270-106), 1 % Pen/Strep/Glutamine (GIBCO, #10378-016), 50 ug/ml Geneticin (GIBCO, #10131-027), 15 ug/ml Hygromcyin (Invitrogen, #10687010) and 1 ug/ml Puromycin (GIBCO, #11138-03). Induction of HTT expression was performed with Doxycycline (Clontech, #631311) at a final concentration of 1ug/ml.

## METHOD DETAILS

### Antibodies

The following antibodies were used: anti-FLAG M2 (Sigma-Aldrich F3165), anti-HTT N18 (Santa Cruz SC-8767), anti-polyQ (Sigma 3B5H10), anti-HAP40 (Santa Cruz SC-69489) and anti-Strep (IBA 2-1507-001).

### Cell lines

B1.21 cells (Huang et al., 2015) are based on HEK293 cells and express FLAG-His tagged human full length 17QHTT upon induction with doxycycline (Dox). B1.21-HAP40 cells (Guo et al., 2018) additionally co-express human HAP40 constitutively. C2.6 cells (Huang et al., 2015) are based on HEK293 cells and express FLAG-His tagged human full length mutant 46QHTT.

HTT128 cells were generated in a similar manner as B1.21 cells. A cDNA coding for 128QHTT with C-terminal fusion to a FLAG-His affinity tag was cloned into the vector pTRE-tight-BI-AcGFP1 (Clontech) for expression of 128QHTT upon induction with Dox. The resulting plasmid pTRE-HTT128Q was verified by restriction analysis and sequencing. HEK293 Tet-ON cells (Clontech) were co-transfected with linearized pTRE-HTT128Q together with a plasmid expressing a hygromycin resistance gene. Positive cell clones were selected by addition of hygromycin to the culture medium. Monoclonal HTT128 cell lines were obtained by limited dilution of positive cell clones.

C.6-HAP40TS and HTT128-HAP40TS cells were generated in a similar manner as B1.21-HAP40 cells. In brief, C2.6 and HTT128 cells were co-transfected with plasmid pBSK/2-CMV-HAP40-TS together with a linearized plasmid expressing a puromycin resistance gene. Positive cell clones were selected by addition of puromycin to the culture medium. The resulting stable cell lines (C2.6-HAP40TS and HTT128-HAP40TS) express, in a constitutive manner, HAP40 C-terminally fused to a Strep affinity tag and mutant HTT (46QHTT and 128QHTT, respectively) upon induction with Dox. These cell lines were used for production of mutant 46QHTT-HAP40 and 128QHTT-HAP40 complexes, respectively.

All generated cell lines were tested negative for mycoplasma by PCR. Induction of mutant HTT expression by Dox was confirmed by Western blot analysis.

### Purification of 17QHTT-HAP40, 46QHTT-HAP40 and 128QHTT-HAP40 complexes

The HTT-HAP40 complexes were purified as described (Guo et al., 2018). In brief, 1.2×10^9^ B1.21-HAP40TS cells, C2.6-HAP40 cells or HTT128-HAP40 cells were harvested 72 h after induction with Dox by centrifugation at 400 g for 10 min. Cells were lysed in 25 mM HEPES, 300 mM NaCl, 0.5 % Tween 20, protease inhibitor, pH 8.0, by rotation at 4° C for 30 min followed by centrifugation of the cell lysate at 30,000 g and clearance by filtration through a 0.2 μm filter. The filtrate was incubated with Strep beads (IBA) for 2-3 h at 4 °C. After washing three times with 25 mM HEPES, 300 mM NaCl, 0.02% Tween 20, pH 8.0, bound proteins were eluted with 25 mM HEPES, 300 mM NaCl, 0.02% Tween 20, 2.5 mM Desthiobiotin, pH 8.0. The eluate was concentrated using Amicon filters.

The HTT-HAP40 complex was further purified by size exclusion chromatography (SEC) using a Superose 6 10/300 increase column (GE Healthcare) in running buffer 25 mM HEPES, 300 mM NaCl, 0.1% CHAPS and 1 mM DTT, pH 8.0. HTT-HAP40 eluted in one narrow-based peak and was concentrated with Amicon ultra 100 kDa filters (Millipore). The purity of purified HTT-HAP40 complex was analysed by SDS-PAGE and silver Coomassie-Blue staining. The integrity of HTT and HAP40 in the HTT-HAP40 complex was confirmed by Western blot analysis.

### Protein thermostability measurement by differential scanning fluorimetry (DSF)

The thermostability of HTT-HAP40 complexes was assessed by DSF (Vedadi et al., 2006). Protein unfolding was monitored by the increase in the fluorescence of SYPRO Orange (Invitrogen). Prior to use, a 100 mM stock of the dye (stored at -20°C) was diluted 1:20 in DMSO and directly added to the sample to a final concentration of 125 µM. The proteins were diluted in sample buffer (25 mM HEPES, 300 mM NaCl, 0.1% CHAPS, 1 mM DTT and 10% glycerol) to concentrations of 0.2 mg/ml. The samples were heated up with a ramp rate of 1 °C/min over a temperature range of 15-95 °C using the qPCR System MX 3005 P (Stratagene). Measurements were performed in triplicate. Unfolding transition temperatures (Tm) were automatically determined by the software of the qPCR machine.

### Protein thermostability measurement by nanoDSF

Protein thermostability was further assessed using low volume differential intrinsic tryptophan scanning fluorimetry (nanoDSF) (Alexander et al., 2014). Here, the intrinsic fluorescence of protein thryptophan residues was directly monitored to eliminate artefacts caused by labeling or modifying the protein. The tested proteins were diluted in sample buffer (25 mM HEPES, 300 mM NaCl, 0.1% Chaps, 1 mM DTT and 10% glycerol) to a final concentration of 0.3 mg/ml. 10 μl samples were manually loaded into nanoDSF Grade Standard Capillaries (NanoTemper Technologies) and transferred to a Prometheus NT.48 nanoDSF device (NanoTemper Technologies). Samples were heated with a linear ramp rate of 1 °C/min over a temperature range from 20-95 °C. Unfolding profiles were determined from changes in the emission wavelengths of tryptophan fluorescence at 330 nm, 350 nm and their ratios. Data was analyzed with the Prometheus PR. Control software (NanoTemper Technologies). Measurements were performed in triplicates.

### Cryo-EM sample preparation, data acquisition and image processing

Cryo-EM samples were prepared and processed as described (Guo et al., 2018). Purified HTT-HAP40 complexes were diluted to 0.5 mg/ml with 25 mM HEPES, 300 mM NaCl, 0.025% CHAPS, 1 mM DTT. 4 μl of samples were applied to R2/4 Quantifoil gold grids with suspended monolayer graphene (Graphenea) and vitrified by plunge-freezing into a liquid ethane/propane mixture using Vitrobot Mark IV (FEI). Data collection was performed on a Titan Krios microscope operated at 300 kV and equipped with a Gatan GIF Quantum energy filter. The calibrated magnification was 105,000X in EFTEM mode, corresponding to a pixel size of 1.35 Å. Images were collected by a K2 Summit direct electron camera (Gatan) using counting mode, with a dose rate of 4 electrons/Å^2^/s. Each exposure (8 s exposure time) comprised 16 sub frames, amounting to a total dose of 32 electrons/Å^2^. Data was recorded using SerialEM software (Mastronarde, 2005) and custom macros. Defocus values ranged from -1.4 to -3 μm. 1127 and 1438 movie stacks were collected for 46QHTT-HAP40 and 128QHTT-HAP40 samples respectively.

Raw frame stacks were subjected to beam-induced motion correction using MotionCor2 (Zheng et al., 2017). Most further processing was performed using RELION (Scheres, 2012). The contrast transfer function parameters for each micrograph were determined with CTFFIND4 (Rohou and Grigorieff, 2015), and all micrographs with a resolution limit worse than 4 Å were discarded. Particles were initially picked with Gautomatch (http://www.mrc-lmb.cam.ac.uk/kzhang/Gautomatch/Gautomatch_Brief_Manual.pdf) using a sphere as template, and extracted with a 160-pixel by 160-pixel box. Reference-free 2D class averaging was performed reiteratively, keeping only particles contributing to well-resolved 2D averages. The resulting particle subsets were used for further three-dimensional classification. The classes with identical detailed features were merged for further auto-refinement to produce the final density map with an overall resolution of 3.6 Å and 4.1 Å for 46QHTT-HAP40 and 128QHTT-HAP40, respectively (Figure S 1, Table S 1). The resolution estimation was based on the gold-standard Fourier shell correlation method using the 0.143 criterion (Scheres and Chen, 2012). All density maps were sharpened by applying temperature factor that was estimated using post-processing in RELION. For visualization, the density maps were filtered based on the local resolution determined using half-reconstructions as input maps.

### Model building

Modeling of 46QHTT-HAP40 and 128QHTT-HAP40 complexes was performed in COOT (Emsley and Cowtan, 2004), using 17QHTT-HAP40 (Guo et al., 2018) as starting model. Regions (1-90, 323-342, 403-660, 960-977, 1049-1057, 1103-1120, 1158-1222, 1319-1347, 1372-1418, 1504-1510, 1549-1556, 1714-1728, 1855-1881, 2063-2091, 2325-2347, 2472-2490, 2580-2582, 2627-2660, 2681-2687, 2926-2944 and 3099-3138) of HTT and regions (1-41, 217-257, 300-313 and 365-371) of HAP40 were not built in the final model, as no well-resolved densities were present in the map. Maps refinements were carried out using Phenix.real_space_refine (Adams et al., 2010) against the respective overall map at resolution of 3.6 Å (46QHTT-HAP40) and 4.1 Å (128QHTT-HAP40), with secondary structure and Ramachandran restrains. The final models were validated using MolProbity (Chen et al., 2010).

The 46QHTT-HAP40 and 128QHTT-HAP40 models were aligned to the 17QHTT-HAP40 model before analysis. Pairwise Cα RMSD was calculated using the PyMOL (Schrödinger) script rmsd_b.py (available at http://pldserver1.biochem.queensu.ca/~rlc/work/pymol/). Maps and models were visualized using Chimera (UCSF).

### Chemical crosslinking coupled to mass spectrometry

Protein complexes were crosslinked and measured essentially as described (Leitner et al., 2014). In short, 40 μg of equimolar mixtures of HTT and HAP40 (0.7 µg / µL stored in 25 mM HEPES, pH 7.4, 300 mM NaCl, 0.05 % CHAPS, 1 mM DTT and 10 % glycerol) were crosslinked by addition of H12/D12 disuccinimidyl suberate (DSS) (Creative Molecules) at a final ratio of 2.5 nmol DSS per 1 µg protein for 30 min at 37 °C while shaking at 650 rpm in a Thermomixer (Eppendorf). After quenching by addition of ammonium bicarbonate to a final concentration of 50 mM and incubation for 10 min at 37 °C, samples were evaporated to dryness, dissolved in 8 M urea, reduced with TCEP (final concentration of 2.5 mM), alkylated with iodoacetamid (final concentration of 5 mM) and digested overnight with trypsin (Promega V5113) in 1 M urea at an enzyme-to-substrate ratio of 1:40. Digested peptides were separated from the solution and retained by a solid phase extraction system (SepPak, Waters) and subsequently enriched by SEC on an ÄKTAmicro chromatography system (GE Healthcare) using a Superdex™ Peptide 3.2/300 column (GE Healthcare) at a flow rate of 50 µL/min of the mobile phase (water/acetonitrile/trifluoroacetic acid 70 %/30 %/0.1 %, vol/vol/vol). UV absorption at a wavelength of 215 nm was used for monitoring the separation. The eluent was collected in fractions of 100 µL in a 96-well plate. The three fractions 1.0 – 1.1 mL, 1.1 – 1.2 mL and 1.2 – 1.3 mL were collected, dried and further analyzed by LC-MS/MS.

Sample amounts fractionated by SEC were normalized and re-dissolved in an appropriate volume of MS buffer (acetonitrile/formic acid 5 %/0.1 %, vol/vol) according to their UV signal (at 215 nm) prior to liquid chromatography (LC)-MS/MS analysis on an Orbitrap Fusion Tribrid mass spectrometer (Thermo Scientific).

Data were searched using *xQuest*. Crosslinking was performed in biological triplicates and each sample was additionally measured in technical duplicates. Crosslinks were only considered during structural analysis, if they were identified in at least 2 of 3 biological replicates with deltaS < 0.95, ld score ≥ 25 and at least one with an assigned false discovery rate (FDR) as calculated by *xProphet* below 0.05. Identified crosslinks were visualized by XiNET (Combe et al., 2015).

A list of all identified links can be found in Table S 2.

### LC-MS/MS analysis

Peptides were separated on an EASY-nLC 1200 (Thermo Scientific) system equipped with a C18 column (Acclaim PepMap 100 RSLC, length 15 cm, inner diameter 50 µm, particle size 2 µm, pore size 100 Å, Thermo Scientific). Peptides were eluted at a flow rate of 300 nL/min using a 60 min gradient starting at 94 % solvent A (water/acetonitrile/formic acid 100 %/0 %/0.1 %, vol/vol/vol) and 6 % solvent B (water/acetonitrile/formic acid 20 %/80 %/0.1 %, vol/vol/vol) for 4 min, then increasing the percentage of solvent B to 44 % within 45 min followed by a 1 min step to 100 % B for additional 10 min. The mass spectrometer was operated in data-dependent-mode with dynamic exclusion set to 60 s and a total cycle time of 3 s. Full scan MS spectra were acquired in the Orbitrap (120.000 resolution, 2e5 AGC target, 50 ms maximum injection time). Most intense precursor ions with charge states 3 – 8 and intensities greater than 5e3 were selected for fragmentation using collision induced dissociation with 35 % collision energy. Monoisotopic peak determination was set to peptide and MS/MS spectra were acquired in the linear ion trap (rapid scan rate, 1e4 AGC target).

## Supplementary Figures and Figure Legends

**Figure S1:**
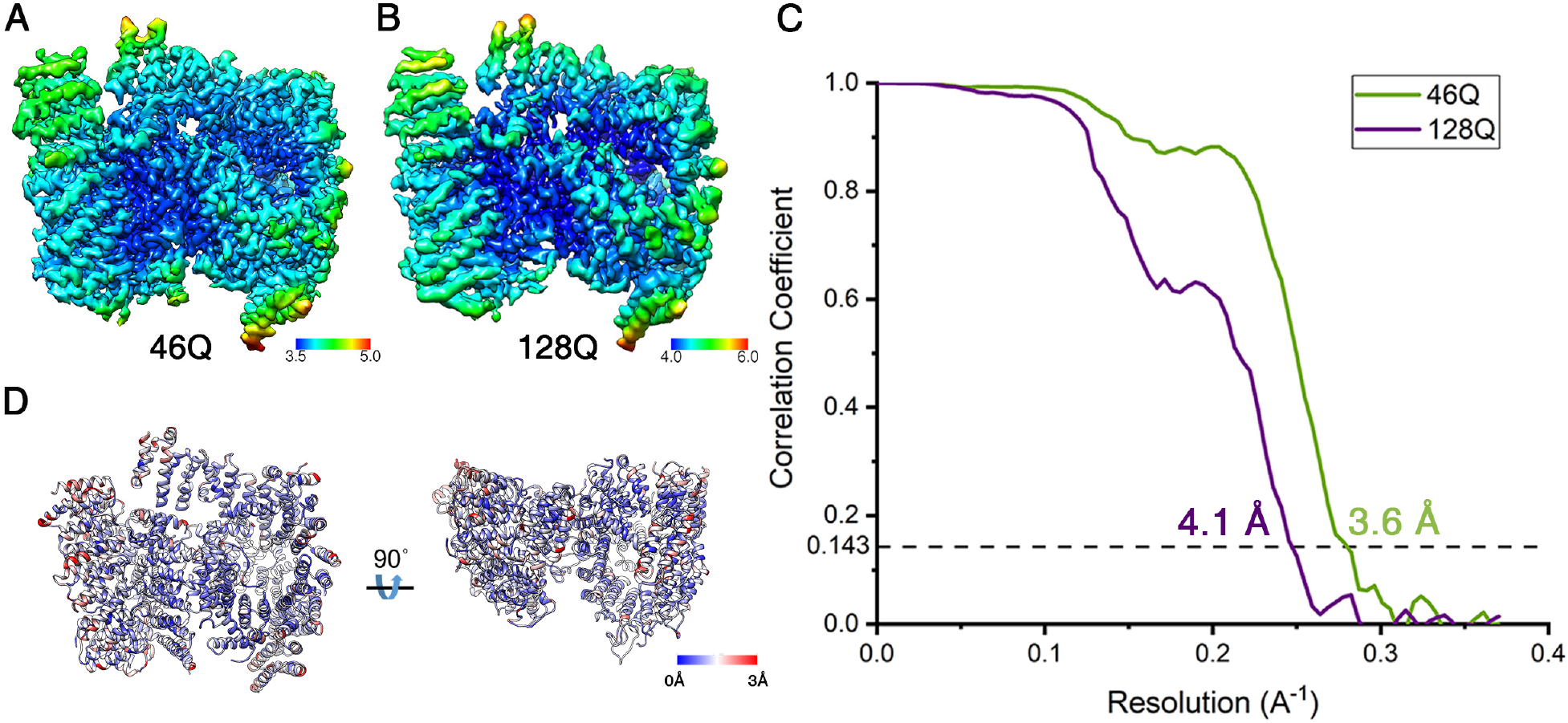
Local resolution of cryo-EM structures. **(A, B)** Density maps of 46QHTT-HAP40 and 128QHTT-HAP40 complexes colored according to local resolution. The maps were filtered according to the local resolution. **(C)** FSC plots of the two structures. The FSC = 0.143 criterion was used for the estimation of the global resolution. Related to Figure 2.

**Figure S2:**
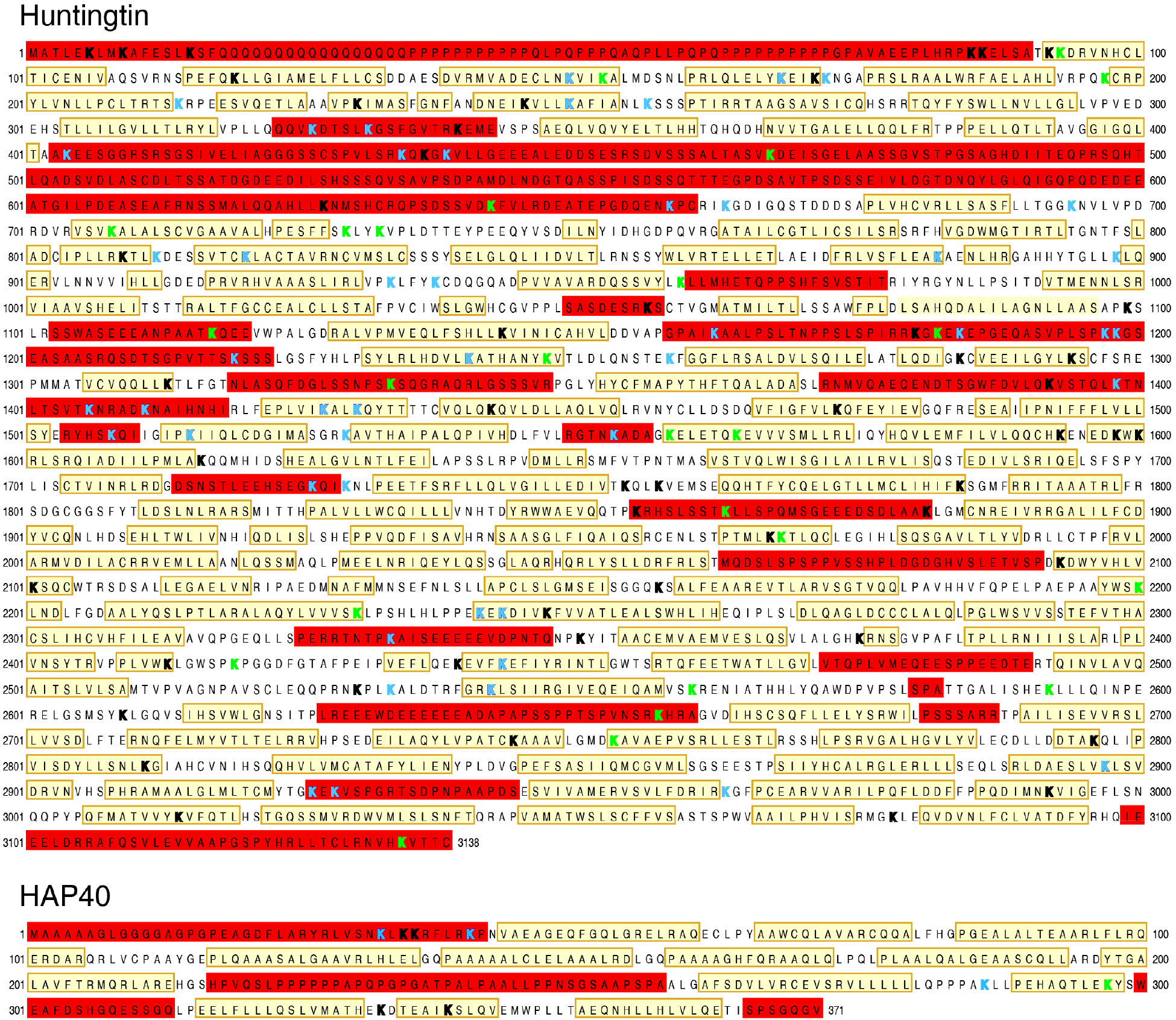
Mapping of monolinks and crosslinks in the sequence of HTT and HAP40. Structural elements of 17QHTT are indicated as follows: invisible in the model (red boxes) and α-helices (yellow boxes). Lysine residues (K) with monolinks (green) or crosslinks (blue) detected in all three HTT-HAP40 complexes (17QHTT-HAP40, 46QHTT-HAP40 and 128QHTT-HAP40) are marked. Only high-confidence crosslinks that were reliably identified in at least two out of three biological replicates are shown (see methods for details). The remaining lysines are marked in black bold. Related to Figure 3.

**Figure S3:**
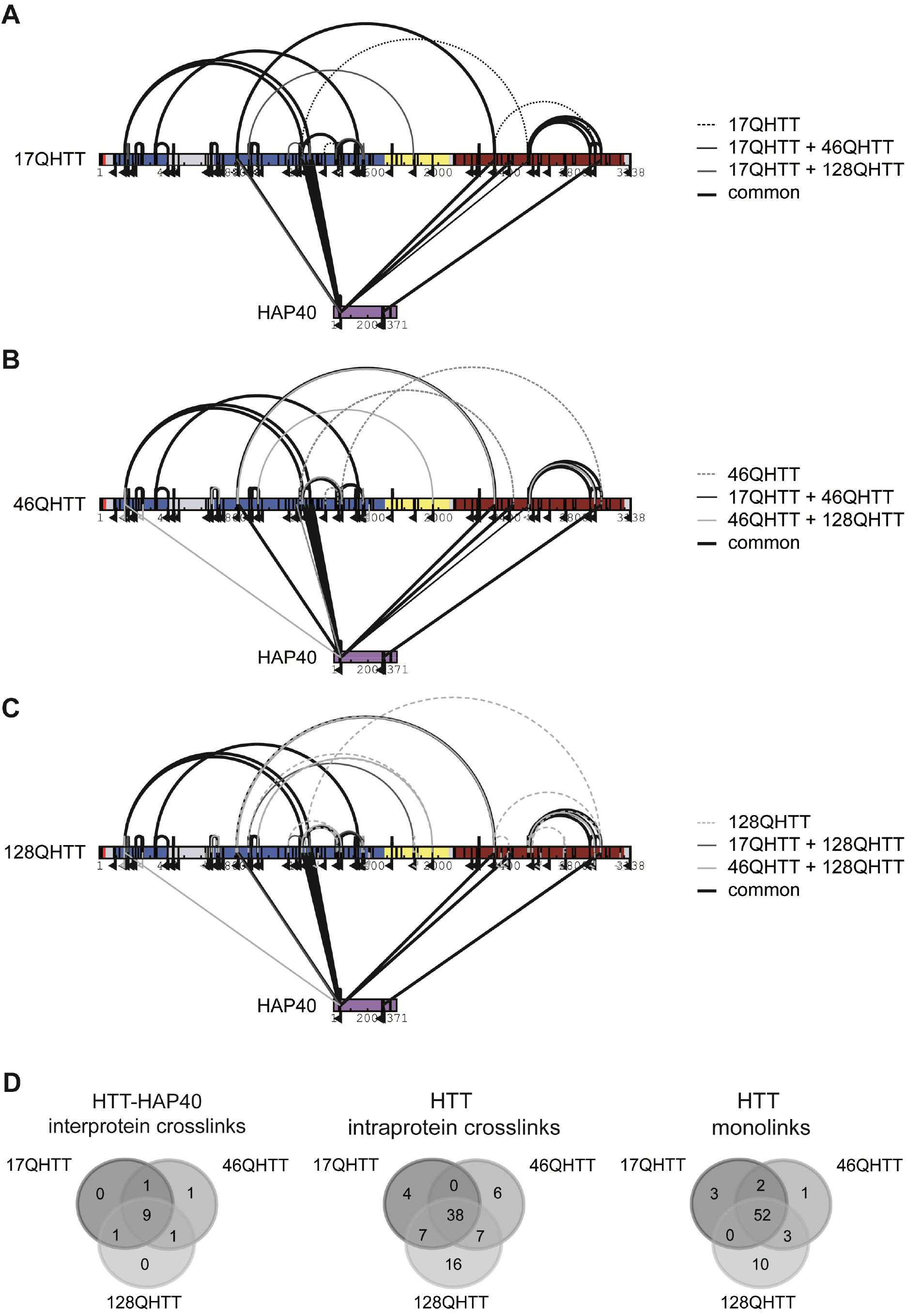
Effects of polyQ expansion on overall crosslink patterns of HTT-HAP40 complexes. **(A, B, C)** Comparative representation of all crosslinks identified in three different **polyQ expanded** HTT-HAP40 complexes, namely 17QHTT-HAP40 (A), 46QHTT-HAP40 (B) and 128QHTT-HAP40 (C). A uniform numbering scheme based on the 17QHTT sequence was used. Crosslinks common to all three complexes are represented by thick, black solid lines. Crosslinks identified in only two of the complexes are represented by thin solid lines (17QHTT and 46QHTT, black; 17QHTT and 128QHTT dark grey; 46QHTT and 128QHTT, light grey). Crosslinks identified in only one of the complexes are depicted by dotted lines (17QHTT, black; 46QHTT, dark grey; 128QHTT light grey). Only high-confidence crosslinks that were reliably identified in at least two out of three biological replicates are shown (see methods for details). Interlinks are shown as straight line, Intralinks as curved line and monolinks as a flag. The HTT N-HEAT domain, bridge domain and C-HEAT domains are colored in blue, yellow and maroon, respectively. HAP40 is colored in purple. (**D**) Venn diagrams showing the overlap of crosslinks identified in the three different HTT-HAP40 complexes (**A, B, C**). Related to Figure 3.

## Supplementary Tables and Table Legends

**Table S1:** Cryo-EM data collection, refinement and validation statistics. Related to Figure 2.

|  | 46Q-Huntingtin-HAP40 complex<br>(EMD-30911)<br>(PDB 7DXJ) | 128Q-Huntingtin-HAP40 complex<br>(EMD-30912)<br>(PDB 7DXK) |
| --- | --- | --- |
| **Data collection and processing** |  |  |
| Magnification | 105,000X | 105,000X |
| Voltage (kV) | 300 | 300 |
| Electron exposure (e-/Å<sup>2</sup>) | 32 | 32 |
| Defocus range (µm) | 1.4-3.0 | 1.4-3.0 |
| Pixel size (Å) | 1.35 | 1.35 |
| Symmetry imposed | C1 | C1 |
| Initial particle images (no.) | 581'397 | 463'213 |
| Final particle images (no.) | 192'473 | 122'992 |
| Map resolution (Å) | 3.6 | 4.1 |
| FSC threshold | 0.143 | 0.143 |
| Map resolution range (Å) | 3.5-5.0 | 4.0-6.0 |
| **Refinement** |  |  |
| Map sharpening *B* factor (Å<sup>2</sup>) | -140 | -187 |
| Model composition |  |  |
| Non-hydrogen atoms | 20491 | 20491 |
| Protein residues | 2.621 | 2.621 |
| R.m.s. deviations |  |  |
| Bond lengths (Å) | 0.0076 | 0.0074 |
| Bond angles (°) | 1.38 | 1.30 |
| Validation |  |  |
| MolProbity score | 1.66 | 1.70 |
| Clashscore | 4.80 | 5.93 |
| Poor rotamers (%) | 0.97 | 0.48 |
| Ramachandran plot |  |  |
| Favored (%) | 93.86 | 94.60 |
| Allowed (%) | 5.90 | 5.24 |
| Disallowed (%) | 0.23 | 0.16 |

**Table S 2: Crosslinking data**. Related to Figure 3.

